# Classical Myelo-Proliferative Neoplasms emergence and development based on real life incidence and mathematical modeling

**DOI:** 10.1101/2025.05.14.654143

**Authors:** Ana Fernández Baranda, Vincent Bansaye, Evelyne Lauret, Morgane Mounier, Valérie Ugo, Sylvie Méléard, Stéphane Giraudier

**Affiliations:** CMAP, CNRS, Ecole polytechnique, Institut Polytechnique de Paris, 91128, Palaiseau, France; Université de Paris, Institut Cochin, INSERM U1016, CNRS UMR8104, F-75014 PARIS, France; Malignant Hemopathies Registry of cote d’Or, INSERM U1231, CHU Dijon Bourgogne, Dijon, France; Univ Angers, Nantes Université, CHU Angers, INSERM, CNRS, CRCI2NA, F-49000 Angers, France; Institut universitaire de France; Université Paris-Cité, Hôpital Saint Louis, INSERM U1131, F-75010 Paris, France

## Abstract

Mathematical modelling allows us to better understand the emergence and evolution of myeloproliferative neoplasms. We tested different mathematical models on a first cohort (patients) (Côte d’Or Registry) to determine the onset and evolution times before JAK2V617F classical myeloproliferative disorders (polycythemia vera and essential thrombocythemia) are diagnosed. We considered the time to diagnosis as the sum of two periods: the time (from embryonic development) for the JAK2V617F mutation to appear, not disappear and enter proliferation, and a second period corresponding to the expansion of the clonal population until diagnosis.

Using increasingly complex models, we show that the rate of active mutation cannot be constant, but rather increases exponentially with age, following the well-known Gompertz model. We found that it takes an average of 63.1 +/- 13 years for the first tumor cell to appear and start proliferating. On the other hand, the expansion time is constant: 8.8 years once the mutation has occurred. These results were validated in an external cohort (national FIMBANK cohort). Using this model, we analyzed JAK2V167F Essential Thrombocythemia versus Polycythemia Vera and found that the time to active mutation in PV is about 1.5 years longer than in ET, while the expansion time is similar.

In conclusion, our multi-step approach and the final age-dependent model for the onset and development of MPN shows that the onset of a JAKV617F mutation should be linked to an ageing mechanism and indicates a period of 8-9 years for the development of a full MPN.

## INTRODUCTION

Myeloproliferative neoplasms (MPN) are chronic disorders issued from the transformation of hematopoietic stem cell (HSC) and related to the acquisition of JAK2 signaling deregulation. The most frequent mutation is JAK2V617F, followed by JAK2 exon12, CALR and MPL mutations. Essential Thrombocythemia (ET) and Polycythemia Vera (PV) are two main MPN that present the JAK2V617F mutation in most cases. ET is characterized by an increased platelet number and PV by (at least) erythroid compartment enlargement. When the JAK2 mutation is responsible for the pathology development, the mutation is present in one allele for most cells for ET, but needs a homologous recombination of JAK2V617F for PV (1).

A third pathology characterized by JAK2 mutation is myelofibrosis, which is often coupled with secondary events such as TET2 or ASXL1 mutations suggesting that this pathology is only driven by JAK2 signaling abnormalities.

To develop ET or PV, two steps are needed: the acquisition of a mutation at HSC level and the development of the cancerous population. Mathematical modelling and statistical analysis of patients’ data can help us better understand the dynamics of mutation emergence and expansion. Using real life registries of MPN and the age of pathology detection in patients, we studied the emergence and expansion of PV and ET in patients with JAK2V617F mutation.

Two types of data can be found in the literature to study the emergence of mutations in HSCs: sequencing data and registries of age of detection. Sequencing data allows to back-track mutations through mathematical models to define the timeline to driver mutation acquisition (2–4). Detection ages allow to model mutation emergence and expansion as done in (5). While we also consider registries of age of detection, once the mutation acquisition time was defined, we studied the time to diagnosis, unlike (5) where the authors studied the behavior of the size of the mutant population. More importantly, all these studies about MPN emergence, have assume a constant mutation occurrence throughout an individual life. Yet, the available data (**Figure 1**) shows a high number of diagnosed patients at older ages which seems inconsistent with this assumption. In fact, a constant mutation occurrence rate would imply that most cases occur at younger ages, when the registries show very few cases.

**Figure 1:**
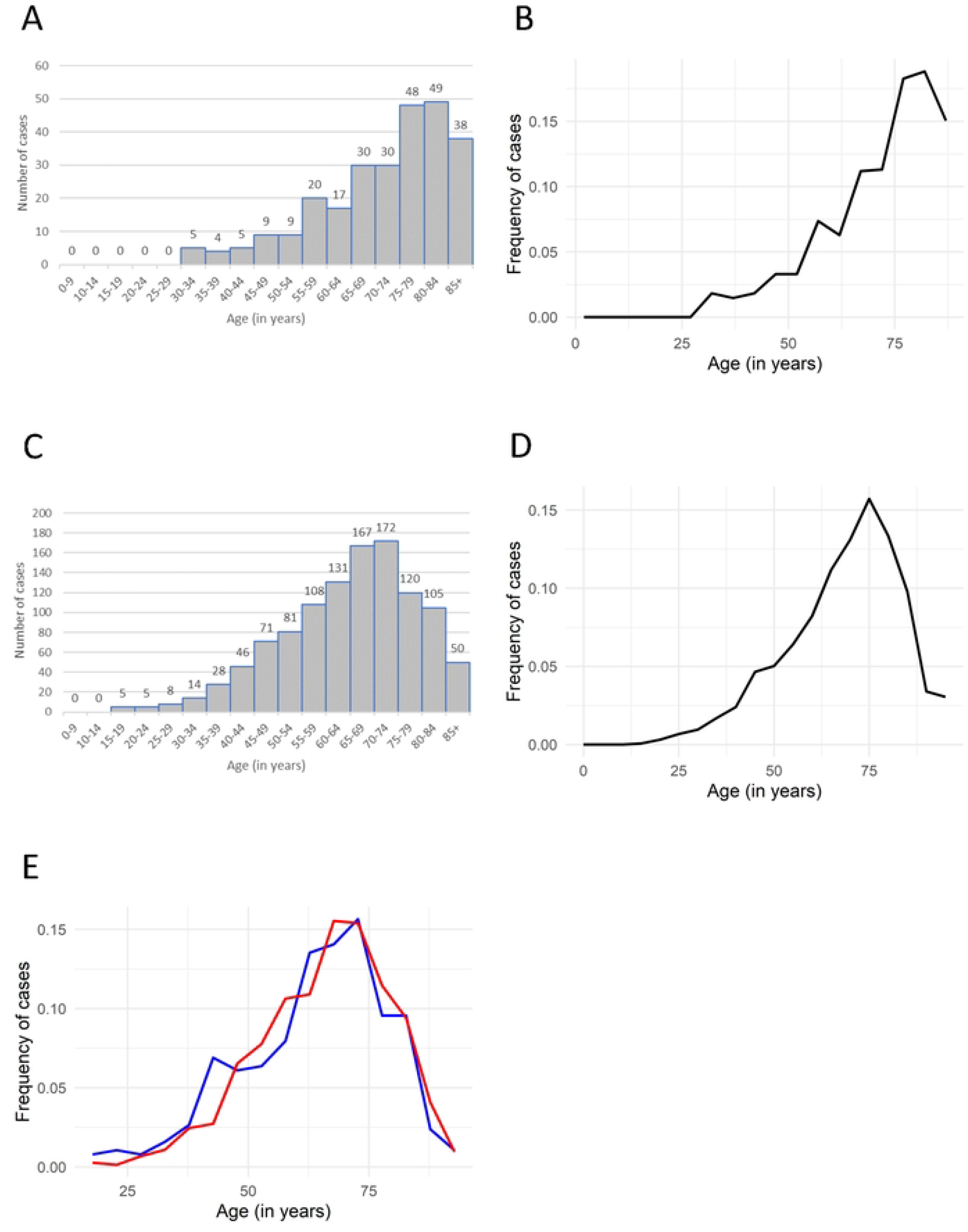
Myeloproliferative cohort descriptions. **A**: Incidence by age of JAK2V617F Classical MPN (ET and PV, excluding myelofibrosis) in the Cote d’Or Regional Registry Database. **B**: Incidence by age of JAK2V617F classical MPN (ET and PV, excluding myelofibrosis) in the Cote d’Or Regional Registry Database after adjustment excluding other causes of death before the MPN diagnosis. **C**: Incidence by age of JAK2V617F Classical MPN (ET and PV, excluding myelofibrosis) in the National BCB-FIMBANK Cohort. **D**: Incidence by age of JAK2V617F classical MPN (ET and PV, excluding myelofibrosis) in the National BCB-FIMBANK Cohort after adjustment excluding other causes of death before the MPN diagnosis. **E**: Frequencies of JAK2V617F ET (red line) and JAK2V617F PV (blue line) by age at diagnosis in the National BCB-FIMBANK Cohort.

Here we defined time to diagnosis as composed of two elements: time for a first cell harboring the JAK2V617F mutation to emerge and enter the cell cycle (T_1_, time of active mutation emergence) and time for cancer population to grow until diagnosis (T_2_, time between emergence and detection). We tested different models of T_1_ and T_2_ to better define MPN development. The models studied in this work for the age of detection of MPN can be divided into two categories: These in which the active mutation rate is constant as previously seen in literature, and these in which it depends on the age of the individual. The first models were analyzed under different assumptions, and we found that they do not appropriately explain (from the statistical point of view) the available data. The second type of models, based on the assumption that the active mutation occurrence increases with age, suggested a much better fit to the data. Hence, to properly explain two available MPN registries, the active mutation occurrence rate rather than being constant, increases exponentially with age (following the well-known Gompertz distribution) conversely to models previously developed.

## RESULTS

### Incidence and frequencies of MPN according to age

Using the cancer registry of the Côte d’Or region in France from 1990 to 2020, we analyzed the ages of detection of 264 patients with different types of MPN according to the type of driver mutation for each pathology (**Table 1**). In this region, life expectancy (and then relative incidence of pathology) can be analyzed. We centered our study in patients diagnosed with either PV or ET presenting the JAK2V617F mutation.

**Table 1:** Number of cases for each age group for the different diseases from the Cote d’Or Regional Registry Database.

| Age | PV JAK2+ | ET JAK2+ | Total JAK2+ cases |
| --- | --- | --- | --- |
| 0 to 4 | 0 | 0 | 0 |
| 5 to 9 | 0 | 0 | 0 |
| 10 to 14 | 0 | 0 | 0 |
| 15 to 19 | 0 | 0 | 0 |
| 20 to 24 | 0 | 0 | 0 |
| 25 to 29 | 0 | 0 | 0 |
| 30 to 34 | 1 | 4 | 5 |
| 35 to 39 | 0 | 4 | 4 |
| 40 to 44 | 1 | 4 | 5 |
| 45 to 49 | 5 | 4 | 9 |
| 50 to 54 | 2 | 7 | 9 |
| 55 to 59 | 8 | 11 | 19 |
| 60 to 64 | 10 | 6 | 16 |
| 65 to 69 | 13 | 15 | 28 |
| 70 to 74 | 12 | 14 | 26 |
| 75 to 79 | 14 | 29 | 43 |
| 80 to 84 | 15 | 30 | 45 |
| More than 85 | 11 | 23 | 34 |
| Total number | 92 | 151 | 243 |

We started by adjusting the data to take into consideration patients that could have died before being diagnosed and thus are not present in the data set (see Supplementary). **Figure 1A** illustrates the histogram of the total cases by ages and **Figure 1B** the corrected frequencies through a linear interpolation by dividing the frequencies of MPN diagnosis at each age by the probability of survival up to this age, the probability of survival was based on the French and European survival data by age and sex of 2021 (6). The resulting corrected data gives us the age distribution of JAK2V617F for patients who have not yet died of anything else. This data indicates that the frequencies of JAK2V617F MPN development increases dramatically with age. We constructed different models and analyzed their fit to the data.

### Modeling mutation occurrence and mutation expansion

When a diagnosis of MPN (such as PV) is performed, the total tumor mass (erythrocytes) represents at least 25% of the erythroid hematopoietic mass (needed increase in Red Cell Mass to diagnose a polycythemia) (7). This suggests a total cancer mass of at least 750 grams in the blood and probably the same mass in the bone marrow. Based on the cell mass, we can speculate that a minimum of one million cell divisions is needed to create such a MPN. This is in agreement with tumor mass estimated in all myeloid chronic myeloproliferative malignancies as previously assessed in Chronic Myeloid Leukemia (8). Indeed, diagnosis is performed when the tumor size reaches 10^12^ cells and with a mean mass of 2.3 ng per cell, the tumor mass can be estimated in the same order of magnitude as PV, and probably ET as well.

Then, we assume that the number of tumor cells needed to perform a diagnosis of PV or ET is of M = 10^12^, and that diagnosis was performed when this size was reached. We consider that the age of detection, T_M_, is the sum of two different periods, T_1_ and T_2_: the first one being the time from embryonic development for the mutation to occur, not disappear, and enter in proliferation (that we will name “mutation occurrence”) and a second time corresponding to the expansion time, once the first cancer cell begins to proliferate until the time of MPN diagnosis.

To define the first period T_1_, we first assumed that the rate of mutation or activation of an existing mutation at HSC level is constant and equal to τ, then the time between two mutations to appear follows an exponential distribution with parameter τ. This assumption has been taken in most of the genetic analysis backtracking reported to date (2–4). We considered that each of these mutant cells have a probability p to proliferate and then be detected. The number (1-p) corresponds to the frequency of non-proliferating mutant cells (just like clonal hematopoiesis does for a long period of time) or of mutant cells whose lineages disappear before reaching detection size of MPN. Under these assumptions, we prove that T_1_ is an exponential random variable with parameter δ = τp (see Supplementary Information), its law is given by f1(t) = δexp(− δt), where represents the age of an individual.

The time T_2_ corresponds to the time taken for a population of cells to reach detection size of 10^12^ cells. Given the high number of divisions needed to reach this size (10^6^divisions is the minimal cell divisions if we considered that no cell die during the expansion process), variability in division time is averaged and it makes sense to consider that the expansion time T_2_ should be similar from one individual to the other. The law of T_2_ then needs to be unimodal, centering around a certain value as for example, a lognormal law. The simplest case is to consider a null variance, where all individuals have the same expansion time, implying that environmental modifications do not influence the growth of the tumor. Following this reasoning, we started by assuming T_2_ was a fixed value and later studied the case when T_2_ is a lognormal random variable.

The times T_1_ and T_2_ were taken as independent, since the time it takes for a cell to become proliferative should not be related to the time it takes for its population to grow to a certain size. In order to consider that the mutation could have appeared at an embryonic level, as suggested in Hermange et al. (5), T_M_ was considered to start 9 months before birth.

Under these assumptions, the detection age T_M_ is distributed as a shifted exponential law, that is

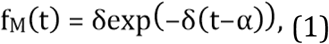

with t corresponding to the age of an individual. The shift α must be a value between 0 and the minimum age of diagnosis in our cohort. We estimated the parameters δ and of the model with a least-squared estimation which minimizes the sum of the square difference between the data and our model for each age. **Figure 2A-2B** illustrates the data of clinical incidence of MPN in real life compared to the estimation, showing a large disparity between the model and the data. The model follows the opposite trend as the data: it gives a strong probability to small ages, while the data is more focused on older ages. It suggests that this model is not appropriate given this type of data. To confirm this, we performed a Chi-square goodness-of-fit test to study the hypothesis that the data follow the distribution of our model and found that it was rejected at a 99 % significance level with a p-value smaller than 10 ^−17^, indicating that a constant occurrence rate together with constant expansion time does not fit with the clinical data.

**Figure 2:**
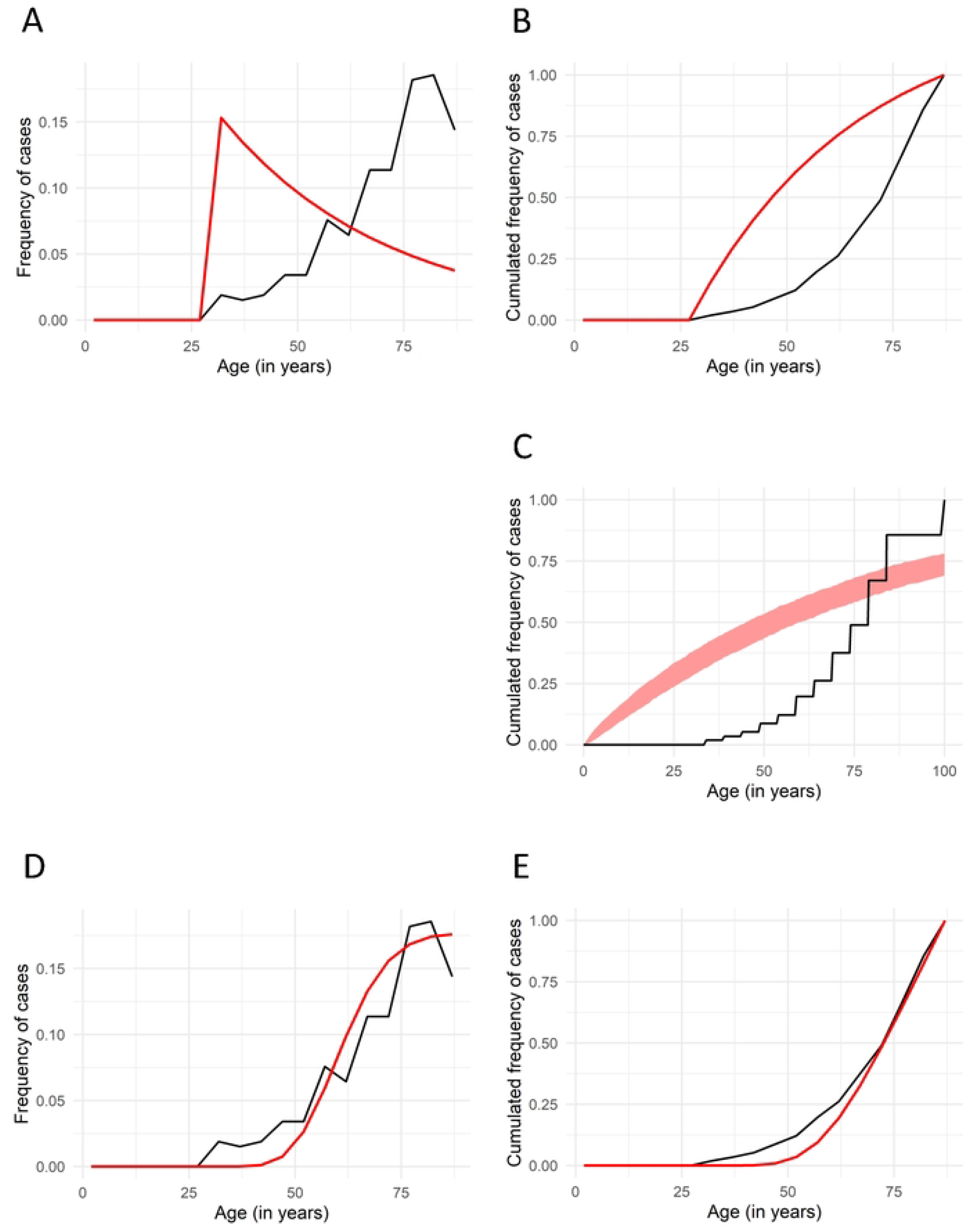
Comparison between data from the Cote d’Or Regional Registry Database and models with constant active mutation rate. **A**: Frequency of cases for the data (in black) and probability density function of the estimation (in red) for the classical model (T_1_ according to constant mutation and mutation activation rates and T_2_ being constant) **B**: Accumulated frequencies of cases for the data from the (black) and cumulative distribution of the estimation (in red) for the classical model. **C**: Accumulated frequencies of cases for the data (in black) and 90% confidence interval of the estimation (in red) for the classical model with individual variability (T_1_ considered as an exponential random variable, with its rate being different from one patient to another, and T_2_ as a constant). **D**: Frequency of cases for the data (in black) and probability density function of the estimation (in red) for the classical model with lognormal MPN growing time (T_1_ according to constant mutation and mutation activation rates and T_2_ taken as a lognormal random variable) **E**: Accumulated frequencies of cases for the data (in black) and cumulative density of the estimation (in red) for the classical model with lognormal MPN growing time.

### Relaxed assumptions on mutation emergence or expansion time

To properly explain the clinical incidence of MPN in the Côte d’Or cohort, we considered two relaxed models that allow additional types of variability: either individual variability on cancer cell emergence or influence of environment on cancer development. The first model considers individual variability (from one patient to another) in the time of mutation occurrence or in the time of mutation development (quiescence/cycling). This takes into account that the biological hypothesis of stress (or environment) influences cancer cell emergence. We considered again T_2_ as a fixed value α and T_1_ as an exponential random variable, but its rate varies from patient to patient. Then, for 1 ≤i ≤N, with N the number of patients, patient i having a personal mutation occurrence rate δ_i_, where the δ_i_’s are considered as a sample of a lognormal(m,s^2^), which distribution is given by

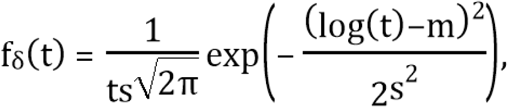

where t represents the age of an individual. This distribution was considered since it gives positive values and allows the δ _i_’s to be centered around a certain value. Then, for patient, the time to mutation emergence and start of proliferation,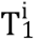 is drawn from an exponential distribution with parameter δ _i_. As in the previous model, we obtain that the law of T_M_ is the shifted exponential distribution (1), but the parameter is different from individual to individual. That is, for individual, the distribution of the age of detection 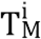 is given by

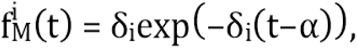

with t being the age of the individual. We can then estimate the values of the parameters m, s and α using the software Monolix (9), a software that is well adapted for this mixed effect modelling. This allows to a simple estimation of the parameters when considering the survival analysis approach based on the SAEM algorithm that gives robust and global convergence. **Figure 2C** illustrates the data compared to the approximation. The estimation presents such a large discrepancy with the data that it is sufficient to reject this model.

A second model considered that external factors may influence the expansion time. Then, the mutation occurrence rate of JAK2V617F is constant while the expansion time of cancer development, T_2_ is taken as a random variable. This could be explained by changes in the invasion process (ie variation in time between two divisions), environmental changes or randomness in the diagnostic process (some patients are diagnosed after vascular events (late detection) while others are diagnosed after a systematic blood cell analysis performed for another pathology (early detection)) or some subjects die (of other unrelated pathologies) before T_2_ was finalized. We assume that T_2_ is a lognormal random variable with parameters (μ, σ^2^), whose distribution is given by

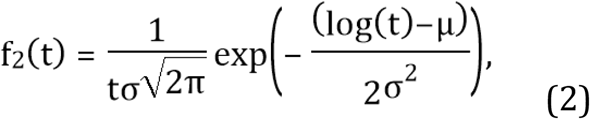

where t represents time after active mutation emergence. Since T_1_ and T_2_ are considered independent, we then have that the law of the age of detection T_M_ is

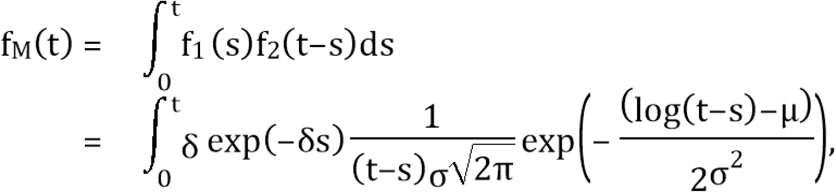

with t the age of an individual.

This model combining an exponential random time for mutation occurrence and a lognormal random time for expansion was also rejected at 99% confidence level through a Chi-square goodness-of-fit test with a p-value smaller than 10^-17^ (**Figure 2D-2E**).

The rejection of both models implies that even when including individual or environmental changes were considered, models with constant active mutation occurrence δ cannot explain the data. Therefore, alternative models based on an increasing probability of JAK2V617F cancer cell emergence with age were mandatory.

### Age-dependency of JAK2V617F mutation emergence

We considered an active mutation occurrence rate that grows with age. Biological motivation for considering an age-dependent mutation occurrence rate δ can be justified by at least three age-dependent modifications of the environment: i-JAK2V617F mutation appearance (from embryo to adulthood), ii- JAK2V617F activation (ie HSC quiescence exit) or iii- the success probability of the expansion after activation.

Let δ(t) be the age-dependent active mutation rate, then, the distribution of T_1_ becomes

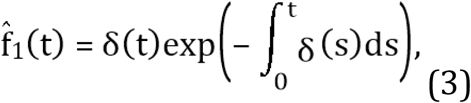

with the age of an individual.

We study two classical models used in survival analysis for age-dependent rates: the Gompertz model and the Weibull model (9). For an age, the respective functions for the mutation occurrence rate are given by

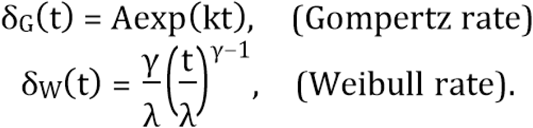

Since the focus is on studying whether age dependence on the active mutation rate can explain the data, we consider the broader model with T_2_ as a lognormal random variable with density function (2). Adding variability to the expansion time T_2_ should give the same or better fit than a fixed value. We later discuss the case of T_2_ constant.

Under these assumptions, the law of T_M_ is given by

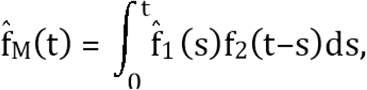

since T_1_ and T_2_ are independent, with 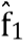 given by (3) and f_2_ by (3) and the age of an individual.

For both the Gompertz and Weibull models, we estimated the parameters using an Expectation-Maximization (EM) algorithm (check Supplementary Information) and performed a Chi-square goodness-of-fit test. We found that the Weibull model was rejected with a p-value smaller than 10^-17^ and the Gompertz model was not rejected with a power of 0.85, indicating that this model seems to properly explain the data.

The Gompertz model has been widely used to model different biological age-dependent rates such as mortality (10) or cancer death rate (11). In our case, it can be explained as follows: we assume that individuals have an initial mutation occurrence rate of A, and an age-related increase in mutation occurrence, represented by a parameter k. Then, for an age t, the active mutation occurrence takes the form

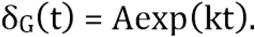

Under this hypothesis, the distribution of the time of active mutation emergence T_1_ is the following

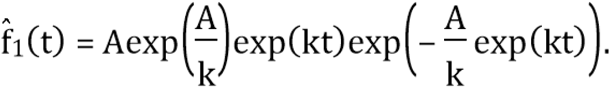

We obtain the following distribution for the age of detection t

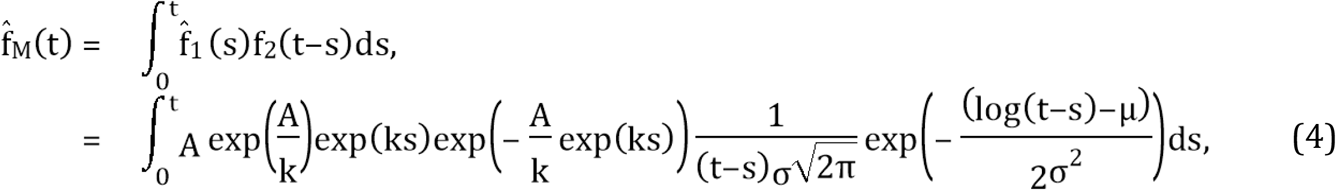

since T_1_ and T_2_ are assumed to be independent.

The numerical results (see Supplementary) show that the variance of T_2_ is quite small, meaning that the expansion time is close to a fixed value.

### Age-dependency of JAK2V617F mutation emergence with different models of tumor growth

We saw that the Gompertz model with lognormal T_2_ was not rejected, but given the small variance of T_2_, we investigated whether a fixed expansion time is sufficient to properly explain the data. When considering T_2_ =α as fixed, we obtained the following distribution for the age of detection t

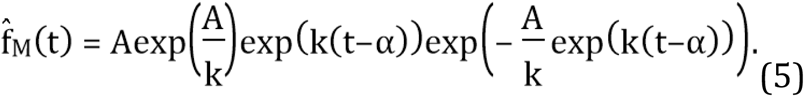

The parameters were estimated using an Expectation-Maximization (EM) algorithm (see Supplementary Information). A Chi-square goodness-of-fit test resulted in not rejecting this model with a power of 0.048. We refer to **Figure 3** for visual comparison between the estimations for fixed T_2_ (**Figure 3A-3B)** and random T_2_ (**Figure 3C-3D)**.

**Figure 3:**
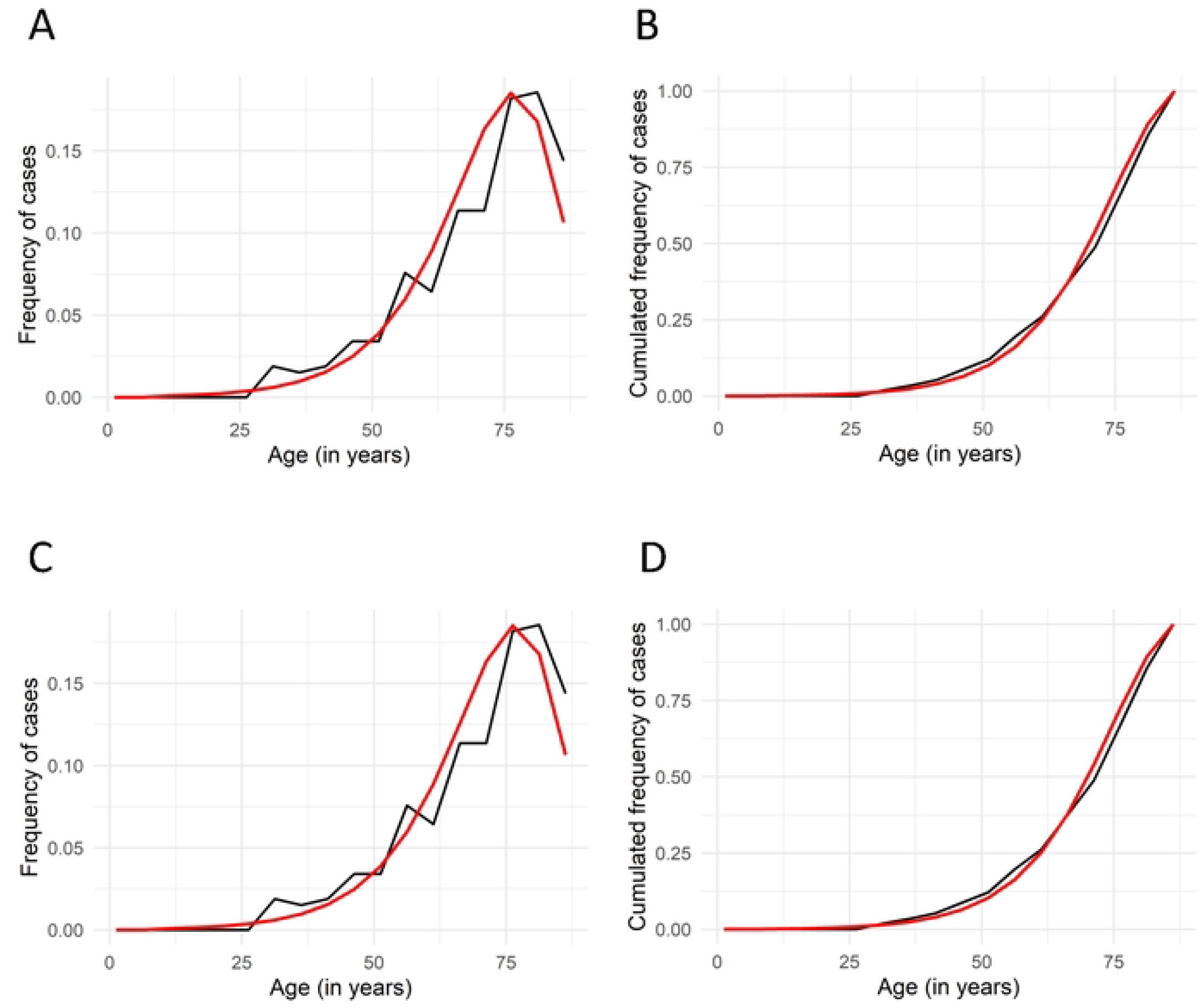
Comparison between data from the Cote d’Or Regional Registry Database and models with aging. **A**: Frequency of cases for the data (in black) and probability density function of the estimation (in red) for the model with aging (T_1_ according to age-dependent mutation and mutation activation rates and T_2_ being constant) **B**: Accumulated frequencies of cases for the data (black) and cumulative distribution of the estimation (in red) for the model with aging **C**: Frequency of cases for the data (in black) and probability density function of the estimation (in red) for the model with aging and lognormal MPN growing time (T_1_ according to age-dependent mutation and mutation activation rates and T_2_ taken as a lognormal random variable) **D**: Accumulated frequencies of cases for the data (black) and cumulative distribution of the estimation (in red) for the model with aging and lognormal MPN growing time.

Since both models are not rejected, we use the Bayesian information criterion (BIC) to select the most appropriate model to explain the data. This criterion measures how well a model can explain the data through the likelihood function, while penalizing the number of parameters in the model to avoid overfitting. A lower BIC is preferred. We obtain that the Gompertz model with constant MPN expansion time (5) (T_2_ as a fixed value) has a BIC of 2079.048 and the Gompertz model with lognormal MPN expansion time (4) has one of 2085.299. This indicates that the fixed expansion time model should be selected. While the inclusion of variability on T_2_ gives a better fit to the data in terms of the likelihood, the gain is small and not compensated by the cost of adding additional unknown parameter. This would imply that, given our data, the Gompertz model with constant MPN expansion time is the most appropriate to explain the data under these assumptions. This suggests that environmental modifications (or other sources of variability) do not have a major impact on the growth phase of the tumor.

By selecting the Gompertz model with fixed expansion time, we found that the expected occurrence time T_1_ is 63 +/- 13 years and the pathology development when the first JAK2V617F stem cell is present, fixed and ready to proliferate is 8.8 years (T_2_) (see Supplementary).

### Validation of the Gompertz model with fixed expansion time in an external cohort

The French group of MPN recently developed a national database on ET and PV (BCB-FIMBANK: French National Cancer Institute (INCa) BCB 2013 and 2022, CHU Angers) (**Table 2**). Given the size of the dataset (n=1111 JAK2V617F MPN patients), we considered this database to be representative of the general population (see **Figure 1C-1D** for the histogram before and after death adjustment), also it is not as exhaustive as the previous database that reported all the cases in the Cote d’Or region. We used this second dataset to validate the Gompertz model with fixed expansion time. We performed an estimation of the parameters through the EM algorithm and then a Chi-Square goodness-of-fit test, which was not rejected with a power of 1 − 2 × 10^−17^ (see **Figure 4A-4B**).

**Table 2:** Number of cases for each age group for the different diseases from the National BCB-FIMBANK Cohort.

| Age | PV JAK2+ | ET JAK2+ | Total JAK2+ cases |
| --- | --- | --- | --- |
| 0 to 4 | 0 | 0 | 0 |
| 5 to 9 | 0 | 0 | 0 |
| 10 to 14 | 0 | 0 | 0 |
| 15 to 19 | 2 | 3 | 5 |
| 20 to 24 | 1 | 4 | 5 |
| 25 to 29 | 5 | 3 | 8 |
| 30 to 34 | 8 | 6 | 14 |
| 35 to 39 | 18 | 10 | 28 |
| 40 to 44 | 20 | 26 | 46 |
| 45 to 49 | 48 | 23 | 71 |
| 50 to 54 | 57 | 24 | 81 |
| 55 to 59 | 78 | 30 | 108 |
| 60 to 64 | 80 | 51 | 131 |
| 65 to 69 | 114 | 53 | 167 |
| 70 to 74 | 113 | 59 | 172 |
| 75 to 79 | 84 | 36 | 120 |
| 80 to 84 | 69 | 36 | 105 |
| 85 to 90 | 30 | 9 | 39 |
| 90 to 94 | 7 | 4 | 11 |
| Total number | 734 | 377 | 1111 |

**Figure 4:**
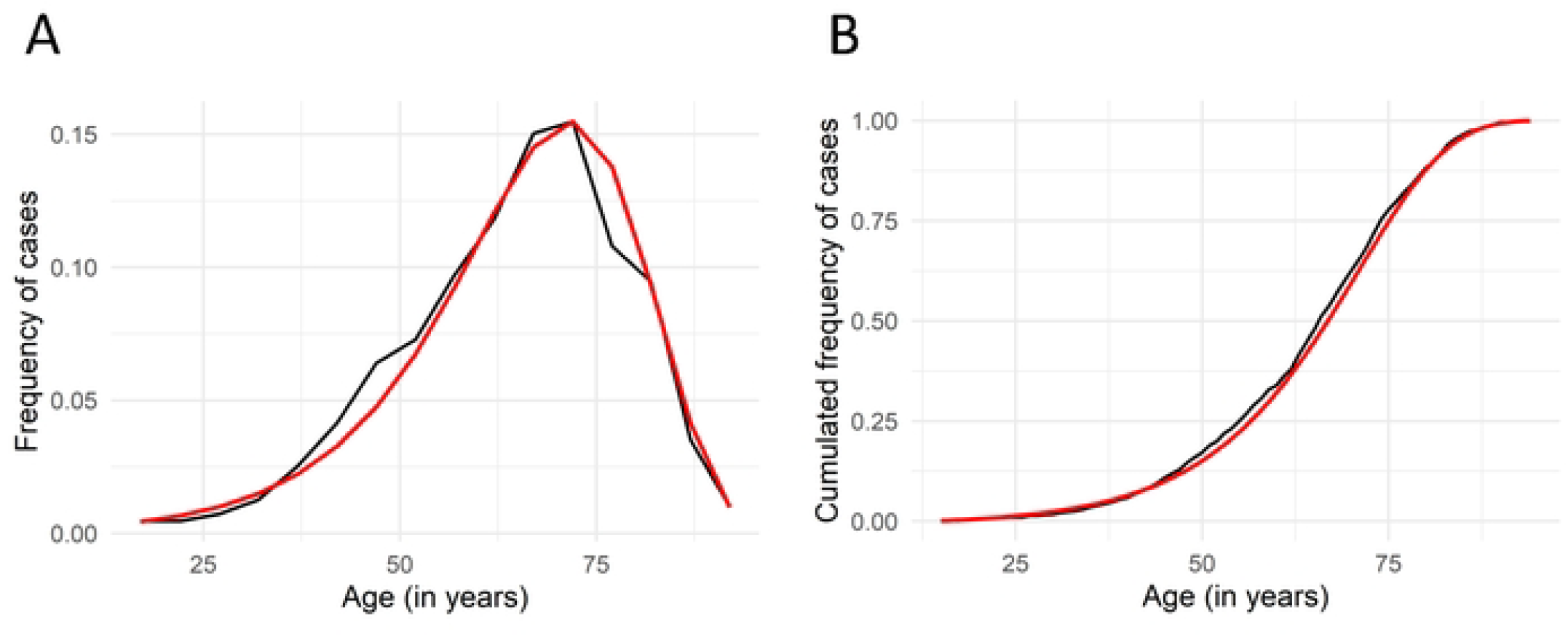
Comparison between data from the National BCB-FIMBANK Cohort and the model with aging. **A**: Frequency of cases for the data (in black) and probability density function of the estimation (in red) for the model with aging (T_1_ according to age-dependent mutation and mutation activation rates and T_2_ being constant) **B**: Accumulated frequencies of cases for the data (black) and cumulative distribution of the estimation (in red) for the model with aging.

This proposed model seems to adequately explain the emergence and proliferation of MPN. This suggests that the mutation rate is variable throughout life and increases exponentially with age according to the Gompertz model. Given these positive results to fit the data, we used the Gompertz model with fixed expansion time to compare the differences in mutation acquisition and expansion between the two pathologies PV and ET.

### Differential analysis of Essential Thrombocytemia and Polycythemia Vera

We hypothesized that ET and PV have a different kinetics because of the double hit in PV compared to single hit in ET: homologous recombination or development of secondary mutations that induces the polycythemia instead of the thrombocytosis only. Given the high number of reported MPN in the FIMBANK cohort, we use this data set to analyze the difference between ET and PV patients.

Looking at their means and standard deviations, PV has a larger mean (65.1 years) by 2 years than ET (63.1 years) (**Figure 1E**) (**Table 3**). The standard deviation of ET is 1.6 years greater (the standard deviation of PV is 13.5 and of ET 15.1), meaning that the ages of detection are spread across a wider range. We first tested whether the ET and PV corrected data came from the same distribution using a Chi-square test, which rejected this hypothesis with a p-value of 10 ^−11^ . Hence, we confirmed that the diagnostic age distributions are different for ET and PV (at a 99% confidence level). We estimated the parameters of each pathology under our model and found that the mean mutation occurrence time (T_1_) is approximately 1.5 years higher for PV which could be explained by the time taken for the mitotic recombination to occur (see Supplementary Information).

**Table 3:**
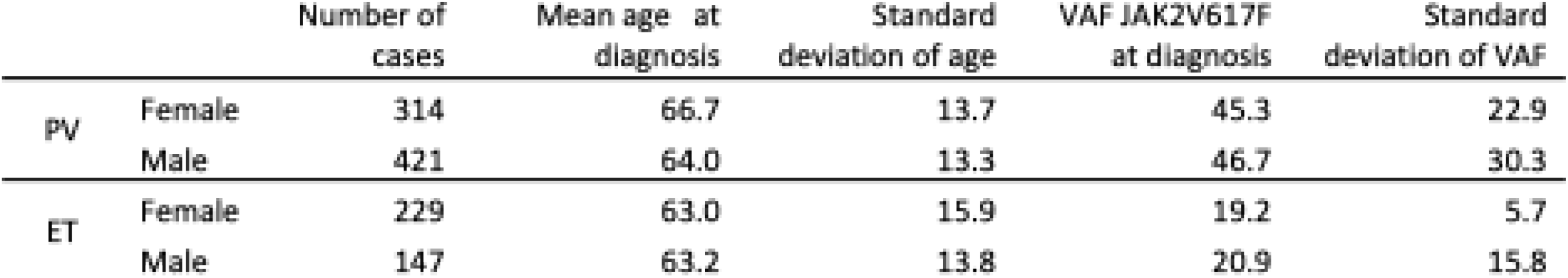
Information on ET and PV from the National BCB-FIMBANK Cohort.

## DISCUSSION

Our present work provides mathematical modeling of JAK2V617F MPN emergence and expansion based on real life incidence and age at diagnosis. This highlights the role of an exponential age-dependent increase in mutation occurrence. Based on the regional exhaustive registry of Côte d’Or (France), and lastly validated in a MPN national registry (BCB-FIMBANK registry), we modelled the occurrence and expansion times of JAK2V617F MPN. Most of the works done so far to decipher when JAK2V617F “first” cell appeared, are based on genetic hierarchy and back-tracking performed on ancient blood samples or bone marrow samples (2–4). These genetic approaches rely upon assumption of a rate (a frequency) of mutation occurrence that is unknown in MPN. It requires old blood or bone marrow samples for genetic analysis (to recapitulate the genetic tree evolution of the original cancer cell) which is obviously very rare and probably biased (12). Genetic analysis inference performed in such approaches are based on mutation occurrence but not necessary on proliferative / oncogenic events. Indeed, it is now well documented that such JAK2V617F mutations could persist for a long time without developing pathologies as seen in clonal hematopoiesis of indeterminate potential (CHIP). Moreover, these genetic back-tracking approaches were based on stable mutation frequencies which overestimate the time of emergence if mutation rate is not constant throughout life. Our approach was based on registry analysis, which differs from classical genetic back-tracking of mutation emergence. We showed that not only the mutation rate cannot be constant, but must grow exponentially with age, following the well-known Gompertz model.

We assumed that two periods of time were required for the development and diagnosis of MPN. The first period (T_1_) corresponds to the mutation acquisition (genetic event from the conception of the embryo) but also its activation (*ie* ensuring its development). This first period can correspond biologically to a cell cycle entry of a long-lasting quiescent cell, but also includes the possibility that a MPN cancer cell disappears due to the cancer cell extinction probability. The second period (T_2_) corresponds to the growth and development of such cancer stem cell until it is diagnosed. This type of modeling has been applied to invasion-fixation of mutants in eco-evolutionary models (13,14) and to the infectious disease propagation (15,16) but less frequently to cancer emergence and development.

Most reported modeling assumes that mutation and invasion rate are aging independent (*ie* a fixed rate of mutation and activation and a fixed probability of emergence of the pathology for a given patient), which would imply that T_1_ follows an exponential law. However, models under this hypothesis show a peak of cases in younger ages, which is inconsistent with the data, where most cases occurred at older ages. We then speculated that myeloproliferative neoplasm emergence could need an age-dependent modeling for T_1_. Age dependence does not necessarily mean that the acquisition of JAK2V617F mutation is higher in aged HSC than in younger ones; it could also be interpreted as a mutation appearing during childhood or even before birth but being activated by an external stimulus long time latter.

The second period (expansion time) is easier to characterize. A model considering the expansion time T_2_ as a fixed value properly fits the data and is preferred to one with T_2_ as a random variable. This suggests that external changes (stress, diet, toxic exposure….) could not significantly alter the behavior of MPN development and that the proliferation time from one “active” cancer cell to the total cancer mass at diagnosis is relatively fixed. This could mean that the most important factor is the proliferative speed related to the mutation acquisition. We then can speculate about the possibility (and risks and benefits) to treat subjects with JAK2V617F clonal hematopoiesis to delay the unset of MPN or prevent the classical thrombotic complications associated to these pathologies.

Our mathematical and statistical study allows us to assess the time of emergence and development of MPN. We found that the average time for a cancer cell to emerge is 63 years and the time from tumor emergence to diagnosis is approximately 8.8 years in line with previous hypothesis (17,18). Based on the Variant Allele Frequency (VAF) of JAK2V617F in the “FIMBANK” cohort, we observe that the mean JAK2V617F VAF in MPN is around 30% (19.9% in ET and 46.1% in PV respectively) and an increased ratio of 1.4% per year (as reported previously in untreated (19) or Hydroxyurea only treated patients (20). This should imply a time of 10 years of time from undetectable VAF to the 30% allele ratio observed at diagnosis in our MPN cohort. This is very close (and slightly higher) to the 8.8 years of tumor expansion time to MPN diagnosis that we calculated in our model. Furthermore, our modeling fits well with the JAK2V617F CHIP which has been described to take approximately a decade to turn into MPN (21–23).

In conclusion, we have used mathematical modelling to demonstrate that the emergence and fixation of JAK2V617F mutation is strongly linked to aging mechanisms and that the expansion time of MPN can be considered as non-random. Taken together, these results highlight the interest in testing all 50-60 years-old inhabitants for JAK2V617F CHIP and question the place of preventive therapies for such subjects.

## References

1. Li J, Kent DG, Godfrey AL, Manning H, Nangalia J, Aziz A, et al. JAK2V617F homozygosity drives a phenotypic switch in myeloproliferative neoplasms, but is insufficient to sustain disease. Blood. 15 mai 2014;123(20):3139–51.

2. Osorio FG, Rosendahl Huber A, Oka R, Verheul M, Patel SH, Hasaart K, et al. Somatic Mutations Reveal Lineage Relationships and Age-Related Mutagenesis in Human Hematopoiesis. Cell Rep. 27 nov 2018;25(9):2308-2316.e4.

3. Van Egeren D, Escabi J, Nguyen M, Liu S, Reilly CR, Patel S, et al. Reconstructing the Lineage Histories and Differentiation Trajectories of Individual Cancer Cells in Myeloproliferative Neoplasms. Cell Stem Cell. 4 mars 2021;28(3):514-523.e9.

4. Mitchell E, Spencer Chapman M, Williams N, Dawson KJ, Mende N, Calderbank EF, et al. Clonal dynamics of haematopoiesis across the human lifespan. Nature. juin 2022;606(7913):343–50.

5. Hermange G, Rakotonirainy A, Bentriou M, Tisserand A, El-Khoury M, Girodon F, et al. Inferring the initiation and development of myeloproliferative neoplasms. Proc Natl Acad Sci U S A. 13 sept 2022;119(37):e2120374119.

6. Ined - Institut national d’études démographiques [Internet]. [cité 21 nov 2024]. Taux de mortalité par sexe et âge - Mortalité, cause de décès – France - Les chiffres. Disponible sur: https://www.ined.fr/fr/tout-savoir-population/chiffres/france/mortalite-cause-deces/taux-mortalite-sexe-age/

7. Hurley PJ. Red cell and plasma volumes in normal adults. J Nucl Med. janv 1975;16(1):46–52.

8. Michor F, Hughes TP, Iwasa Y, Branford S, Shah NP, Sawyers CL, et al. Dynamics of chronic myeloid leukaemia. Nature. 30 juin 2005;435(7046):1267–70.

9. Lavielle M. Mixed Effects Models for the Population Approach: Models, Tasks, Methods and Tools [Internet]. 1re éd. Chapman and Hall/CRC; 2014 [cité 21 nov 2024]. Disponible sur: https://www.perlego.com/book/1663148/mixed-effects-models-for-the-population-approach-models-tasks-methods-and-tools-pdf?utm_source=google&utm_medium=cpc&campaignid=15913700346&adgroupid=165939051780&gad_source=5&gclid=EAIaIQobChMI_8-hl83tiQMVM4toCR30ZhUUEAAYASAAEgLe_fD_BwE

10. Finch CE, Pike MC, Witten M. Slow mortality rate accelerations during aging in some animals approximate that of humans. Science. 24 août 1990;249(4971):902–5.

11. Riffenburgh RH, Johnstone PA. Survival patterns of cancer patients. Cancer. 15 juin 2001;91(12):2469–75.

12. Hirsch P, Mamez AC, Belhocine R, Lapusan S, Tang R, Suner L, et al. Clonal history of a cord blood donor cell leukemia with prenatal somatic JAK2 V617F mutation. Leukemia. août 2016;30(8):1756–9.

13. Champagnat N, Ferrière R, Méléard S. Unifying evolutionary dynamics: From individual stochastic processes to macroscopic models. Theoretical Population Biology. 1 mai 2006;69(3):297–321.

14. Billiard S, Smadi C. The interplay of two mutations in a population of varying size: A stochastic eco-evolutionary model for clonal interference. Stochastic Processes and their Applications. 1 mars 2017;127(3):701–48.

15. Barbour AD, Hamza K, Kaspi H, Klebaner FC. Escape from the Boundary in Markov Population Processes. Advances in Applied Probability. 2015;47(4):1190–211.

16. Barbour A, Reinert G. Approximating the epidemic curve. Electronic Journal of Probability. janv 2013;18(none):1–30.

17. Radivoyevitch T, Hlatky L, Landaw J, Sachs RK. Quantitative modeling of chronic myeloid leukemia: insights from radiobiology. Blood. 10 mai 2012;119(19):4363–71.

18. McKerrell T, Park N, Chi J, Collord G, Moreno T, Ponstingl H, et al. JAK2 V617F hematopoietic clones are present several years prior to MPN diagnosis and follow different expansion kinetics. Blood Adv. 13 juin 2017;1(14):968–71.

19. Stein BL, Williams DM, Wang NY, Rogers O, Isaacs MA, Pemmaraju N, et al. Sex differences in the JAK2 V617F allele burden in chronic myeloproliferative disorders. Haematologica. juill 2010;95(7):1090–7.

20. Kiladjian JJ, Klade C, Georgiev P, Krochmalczyk D, Gercheva-Kyuchukova L, Egyed M, et al. Long-term outcomes of polycythemia vera patients treated with ropeginterferon Alfa-2b. Leukemia. mai 2022;36(5):1408–11.

21. Nielsen C, Birgens HS, Nordestgaard BG, Kjaer L, Bojesen SE. The JAK2 V617F somatic mutation, mortality and cancer risk in the general population. Haematologica. mars 2011;96(3):450–3.

22. Nielsen C, Birgens HS, Nordestgaard BG, Bojesen SE. Diagnostic value of JAK2 V617F somatic mutation for myeloproliferative cancer in 49 488 individuals from the general population. Br J Haematol. janv 2013;160(1):70–9.

23. van Zeventer IA, de Graaf AO, Salzbrunn JB, Nolte IM, Kamphuis P, Dinmohamed A, et al. Evolutionary landscape of clonal hematopoiesis in 3,359 individuals from the general population. Cancer Cell. 12 juin 2023;41(6):1017-1031.e4.

